# An ex vivo human model for safety assessment of immunotoxicity of engineered nanomaterials

**DOI:** 10.1101/2023.06.29.547008

**Authors:** Josephine Blersch, Birgit Kurkowsky, Anja Meyer-Berhorn, Agnieszka K. Grabowska, Eva Feidt, Ellen Junglas, Wera Roth, Dominik Stappert, Armin Kübelbeck, Philip Denner, Eugenio Fava

**Affiliations:** Deutsches Zentrum für Neurodegenerative Erkrankungen e.V. (DZNE), CRFS-LAT, Venusberg-Campus 1/99, 53127 Bonn, Germany; Deutsches Zentrum für Neurodegenerative Erkrankungen e.V. (DZNE), CRFS, Venusberg-Campus 1/99, 53127 Bonn, Germany; Ruprecht Karl University of Heidelberg, IPMB - Institute of Pharmacy and Molecular Biotechnology, SILVACX, Im Neuenheimer Feld 329, 69120 Heidelberg, Germany; SILVACX, Life Science Inkubator Betriebs GmbH & Co. KG, Ludwig-Erhard-Alle 2, 53175 Bonn, Germany

**Keywords:** nanosafety, nanotoxicology, nanomaterials risk assessment, immunotoxicology, high-throughput screening, high-content screening, NLRP3 inflammasome, immune response, PBMCs, safety by design

## Abstract

The unique physicochemical properties of nanomaterials (NM) and engineered nanomaterials (ENM) have pushed their use in many applications ranging from medicine to the food industry, textiles, and many more fields. Thus, human exposure to NM and ENM is growing by the day. However, the current toxicity tests do not reflect the special characteristics of ENM and are not developed for ENM risk assessment. Here we propose a high-throughput cell-based assay using human peripheral blood mononuclear cells (PBMCs) that can monitor the effects of NM and ENM on cytotoxicity and innate immunity. The proposed assay is fully automated and miniaturized, with excellent assay performance parameters (Z’-score >0.5), amenable for large screening campaigns in industrial setting. Immunotoxicity data for ENM safety assessment are collected in dose-response format. At different states, multiparametric readouts for cytotoxicity, and innate immunity are conducted in a combinatorial method, avoiding ENM-induced bias by endotoxin contamination. Integrating this high-dimensional data, allows (i) holistic safety assessment of immunotoxicity effects caused by ENM, classifying safe and toxic ENM phenotypes, and (ii) to deconvolve mode of action of the ENM effect on the PBMCs. As added value the data obtained can be used to troubleshoot ENM or for a safe-by-design approach in product development.

## 1. Introduction

The use of engineered nanomaterials (ENM) in numerous industry sectors contributes directly to a parallel increase in environmental and human exposure to ENM. ENM, with sizes below 100 nm ^[1]^ are abundant in food, cosmetics, and medicine and can be manufactured from a vast assortment of materials with little consistent oversight or regulation.

The success of the use of ENM is mainly due to their unique characteristics—among them, physicochemical, mechanical, electrical, and optical properties—that diverge significantly from their corresponding non-nano counterparts. At the same time, due to these novel characteristics, there is a remarkable gap in the understanding of the environmental, ecological, and health risks of ENM. Indeed, the majority of assays available for testing ENM safety and toxicity have been developed for small (i.e., non-nano) molecules, and they may be limited in the assessment of ENM toxicity due to their insufficient sensitivity to the unique ENM properties ^[2–4]^. A consequence of these limitations is that the toxic effects of many ENM have not been thoroughly characterized; hence, we have only limited knowledge of their safety. Therefore, the scientific community and regulatory bodies have advocated for the development of novel assays specifically designed for ENM, which would have a closer relation to humans, be amenable to high-throughput screening (HTS), and provide a mechanistic view of the processes ^[5-7]^.

Among many aspects of ENM safety assessment, the inflammatory response is a critical one, as it is involved in the majority of the affected organs, including the lungs, skin, and bowel. One of the first responses of the human body to ENM is the triggering of the inflammatory response elicited through the activation of the innate immune system. Upon recognition of ENM from the innate immune system, cell stress or damage and pro-inflammatory responses are induced, leading to a systemic inflammatory response, cytokine storm, or in case of continuous activation, to chronic diseases such as inflammatory bowel disease, rheumatoid arthritis or neuroinflammatory and neurodegenerative diseases ^[8, 9]^. Inflammasome activation, which is one type of innate immune activation, occurs in a two-stage process, consisting of (i) priming and (ii) activation ^[10]^. The first step, priming of naive (i.e., unprimed) cells, is initiated through the recognition of pathogen-associated molecular patterns (PAMPs), host-derived danger-associated molecular patterns (DAMPs), and nanoparticle-associated molecular patterns (NAMPs, e.g., nanomaterials but also protein aggregates) ^[11]^ through pattern recognition receptors (PRRs) such as Toll-like receptors (TLRs) on cells of the innate immune system as for instance macrophages and monocytes. In brief, upon TLR activation (e.g., by lipopolysaccharides (LPS) from the cell membrane of gram-negative bacteria ^[12, 13]^), kinase-mediated intracellular signaling cascades become activated, leading to the translocation of transcription factor NF-kB from the cytoplasm to the nucleus. This initiates the production of the pro-inflammatory cytokines tumor necrosis factor-α (TNFα), pro-interleukin 18 (pro-IL18), and pro-interleukin-1ß (pro-IL1ß). Further, components of the inflammasome are expressed, for instance, NACHT, LRR, and PYD domains-containing protein 3 (NLRP3) ^[14, 15]^. NLRP3 belongs to the NOD-Like Receptor (NLR) inflammasome family containing a nucleotide-binding domain (NOD) and a C-terminal leucine-rich repeat (LRR) receptor. In summary, this first step induces a “primed” state of the cells, which is marked by the intracellular expression of pro-IL1ß and TNFα. The second step leads to the activation of the inflammasome. There are at least three routes leading to inflammasome activation: (i) the lysosomal pathway is activated through particulate matter such as nanomaterials, marked through cathepsin release from ruptured lysosomes; (ii) intrinsic danger signals such as ATP (engaging P2X7 receptor) or microbial toxins, like nigericin inducing K^+^ efflux and Ca^2+^ influx, trigger inflammasome assembly; or (iii) activation through reactive oxygen species (ROS) ^[10]^. Inflammasome activation leads to the assembly of a multiprotein complex, which in turn leads to the maturation and release of IL1ß and IL18 facilitated through caspase-1, which is recruited as pro-caspase-1 to the inflammasome ^[16]^. Inflammasome activation induces an inflammatory apoptosis, called pyroptosis, resulting in a lytic cell death ^[17]^.

ENM can act at both stages of innate immune activation. ENM can give the priming signal to the cells (i) by direct binding to TLRs, or (ii) indirectly through the release of ROS-induced HMGB1 (high mobility group box 1) and activation of TLR-4, as in the case of amorphous silica ENM on human umbilical vein endothelial (HUVEC) cells ^[18]^. ENM are also shown to have unintentional priming effects in primary human monocytes, activating TLR-4 through endotoxin contamination on ENM ^[19, 20]^. Contamination with endotoxins can occur at any stage during synthesis and handling. Specially ENM with cationic and hydrophobic surfaces are prone to binding endotoxins through negatively charged phosphate groups or binding the lipid structure through hydrophobic-hydrophilic interactions. Between 40% ^[20]^ and 60% ^[19]^ of ENM formulations were found contaminated with endotoxins (levels > 1.0 EU/ml). This endotoxin contamination can serve as a priming signal, while ENM themselves can act as an activation signal. This means that through undetected endotoxin contamination, potentially half of the generated immunotoxicity and immune activation data could be biased due to misinterpretation of experimental results.

Looking at the activation signal alone, several types of ENM were shown to induce NLRP3 inflammasome activation subsequent to the priming signal. Among them are (i) fibrous structures such as asbestos fiber ^[21]^ and monosodium urate crystals (MSU) ^[22]^ from nano-to micro range; (ii) metal oxides such as silica (SiO_2_) ^[18, 23-26]^, TiO_2_^[24, 27]^, and CeO_2_^[28]^; (iii) carbo-nanotubes ^[29]^; (iv) metals such as silver ^[30]^; and (v) ENM of organic materials such as polystyrene ENM ^[31]^. SiO_2_ NM were shown to activate the NLRP3 inflammasome leading to Caspase-1-dependent IL-1ß release in a size-^[25]^ and shape-dependent manner ^[28, 32]^. In summary, for inflammasome activation, the particulate characteristic of ENM itself plays a role as much as the physicochemical properties of ENM.

For the safe use of ENM, it is thereby essential to test the materials in an early phase of product development for effects on the immune system to prevent the discovery of adverse events in later stages. This especially applies to effects on the innate immune system, which is the first line of the human body’s defense system. The panel of specific nanomaterial testing guidelines (OECD TG 407, 412, 413, ISO 10093-4, ICH S8) does not include inflammasome activation for immunotoxicity assessment. Still, following the WHO suggestions, to get insight into ENM-caused immune reactions with innate immune activation being the major component, it is necessary to define novel strategies to test ENM immunotoxicity ^[2]^.

Typically, NLRP3 inflammasome activation through ENM is measured through IL-1ß detection in culture supernatants of stimulated cells. Most commonly, LPS-primed, immortalized mouse macrophage ^[21, 32]^ or human leukemia monocyte (THP-1) cell lines ^[33]^, but also human PBMCs ^[26]^ are used. Sharma et al. reported combined cell viability (via MTT cell metabolism assay) and IL1ß release data (determined through enzyme-linked immunosorbent assay (ELISA)) from silica ENM and Alum on naive, NLRP3-, or Caspase-1-knockout THP-1 cells ^[33]^. However, extensive controls are needed in order to ensure ENM do not interfere with the spectrophotometric readout as shown in the results of spectrophotometric and spectrofluorometric assays (e.g., Alamar blue and tetrazolium-based assays, such as MTT) by binding of ENM to the dye. This makes the broad usage of such methods challenging for ENM ^[34]^. Kroll et al. also showed that ENM can interfere with ELISA results by cytokine adsorption to the particle surface and the consequent creation of a non-biological artifact ^[35]^. Therefore, spectrophotometric and spectrofluorometric assays should be avoided or cross-checked with spectrophotometric-independent quantifications. Moyano et al. showed good correlations of gene expression data between *in vitro* and *in vivo* data by comparing mRNA levels of IL6, TNFα, and IFNγ and other cytokines *in vitro* in splenocytes and six hours post intravenous injection into mice ^[36]^. Nevertheless, testing systems based on murine cells inherently conflict with host-specific immune reactions ^[37]^. Additionally, immortalized cell lines cannot properly reflect conditions of human physiology or donor variability. Lastly, although the assessment of released cytokines in isolation gives some insight into the immune system, this alone reveals only a snippet of the final part of the complex pathways involved in the regulation of the immune response; thus, it is not sufficient for a complete understanding of ENM effects on the innate immune system.

Here we describe an approach on the base of an *ex-vivo* human tissue-like system for high-throughput screening of innate immune effects caused by micro- and nanomaterials. The assay combines and correlates multiparametric readouts from high-content imaging with homogenous readouts of culture supernatants. Using this method, functional insights—such as endotoxin contamination of ENM, cytotoxicity, cytokine expression, inflammasome activation, and cytokine release—can be gained from a vast variety of materials under uniform conditions. Here we assess at an early time point the innate immune response that activates signaling cascades, provoking an inflammatory response to ENM and NM exposure.

## 2. Results

To assess, the effects of ENM on the human immune system, we established an *ex vivo* assay based on human peripheral blood mononuclear cells (PBMCs). PBMCs comprise the main cellular components of the innate immune system and have been used extensively in the scientific community as a proxy system to study the immune response ^[38]^. Our assay has been specifically developed and validated for testing and assessing the effect of ENM. The assay also takes into consideration donor variability as we reference each PBMC sample to a proprietary database collecting data from a human population (data not shown). The assay is based on a high-content phenotyping readout that can be combined with a more classical homogenous readout to enhance the depth of the analysis. Furthermore, our assay is fully automated and can be used in a high-throughput fashion to scale the analysis of large numbers of ENM.

The assay is miniaturized to a 384-well plate format, and as in an industrial setting, all steps are set up on an automated platform (e.g., liquid handling pipetting robots, acoustic liquid handling for compound spotting, confocal high-content imaging microscope), allowing high sample throughput while increasing assay robustness and reproducibility. A primary screen was designed in dose-response format to understand the dynamic of ENM-induced effects on the PBMCs. Two states are tested simultaneously: (i) the naive state and (ii) primed state of PBMCs, both treated with different doses of ENM. In the naive state, we can identify (i) cytotoxicity effects of ENM; (ii) ENM-inducing priming effects through the engagement of TLR receptors (by direct targeting or endotoxin contamination); and (iii) ENM-inducing priming in PBMCs and triggering of the inflammasome pathway. Testing different doses of ENM on primed PBMCs allows identifying whether ENM activate the innate immune system. Technically, 14 ENM in 10 doses can be screened in this format on a single 384-well plate, where additional controls such as non-ENM controls, solvent dose ranges, and reference ENM (SiO_2_-OH in 25 nm diameter) dose ranges are located. Those controls serve as internal quality controls and are used to calculate intra- and inter-plate variances. Assay performance parameters such as the coefficient of variation, Z’-score, signal window (SW), and coefficient of variation (CV) were calculated and are summarized in Table 1 and Figure S1. Assay performance measures with Z’-score above 0.5, large signal window, and CVs below 20% for intra- and inter-plate variance are indicators of a robust, rugged, and scalable screening platform. Such measures also serve to emphasize that data of one dose range of ENM are representative. Identified ENM formulations can be tested in a secondary screen for their mode of action using a set of inhibitors regarding the inflammasome pathway or another hypothesized mode of action on a fixed dose of ENM with varying doses of inhibitor or vice versa on different states of the PBMCs.

**Table 1:** Assay performance measures as Z’-score and signal window (SW) for minimum and maximum signal of reference ENM probes (n=5 plates).

| feature | state | Z' | SW | Maximum Signal |  | Minimum Signal |  |
| --- | --- | --- | --- | --- | --- | --- | --- |
|  |  |  |  | mean | sd | mean | sd |
| PI positive cells [%] | naive | 0.90 | 225.25 | 98.82 | 0.34 | 12.74 | 2.47 |
| PI positive cells [%] | primed | 0.87 | 44.93 | 97.44 | 1.71 | 9.52 | 2.05 |
| Activationrate [%] | primed | 0.68 | 24.62 | 70.87 | 1.53 | 15.05 | 4.51 |
| IL1b release [pg/ml] | primed | 0.72 | 11.99 | 7125.56 | 315.44 | 1886.12 | 170.0 |

### ENM and NM induce Cytotoxicity

To understand ENM-induced cytotoxic effects, human PBMCs were incubated with medium for two hours and treated with a dose range of ENM for four hours. Live/dead staining with propidium iodide (PI), counterstained with Hoechst33342 was performed for the final 30 minutes of incubation. Cells were analyzed by high-content imaging (HCI), and the rate of PI-positive cells was calculated relative to the total cell population. To understand the background level of PI-positive cells, the median percentage plus three standard deviations (17%) was calculated from non-ENM treated (naive) controls and can be seen in Figure 2B as a black dotted line. Any measure above that threshold is considered a significant change in PI-positive cells, reflecting cytotoxicity. To understand the effect of ENM decoration on immunotoxicity, five different surface modifications providing different functionalities of silica ENM of 25 nm diameter were tested in this assay. Silica ENM with -OH groups on the surface showed the highest degree of PI-positive cells, followed by –SH and phosphate-modified silica ENM. Interestingly, surface modification with arginine or formulation with PEG completely abolished the general cytotoxic effect of SiO_2_-OH ENM, keeping levels of PI-positive cells below the threshold. This data suggests that the cytotoxicity of 25 nm Silica ENM is dependent on the surface functionalities.

**Figure 1:**
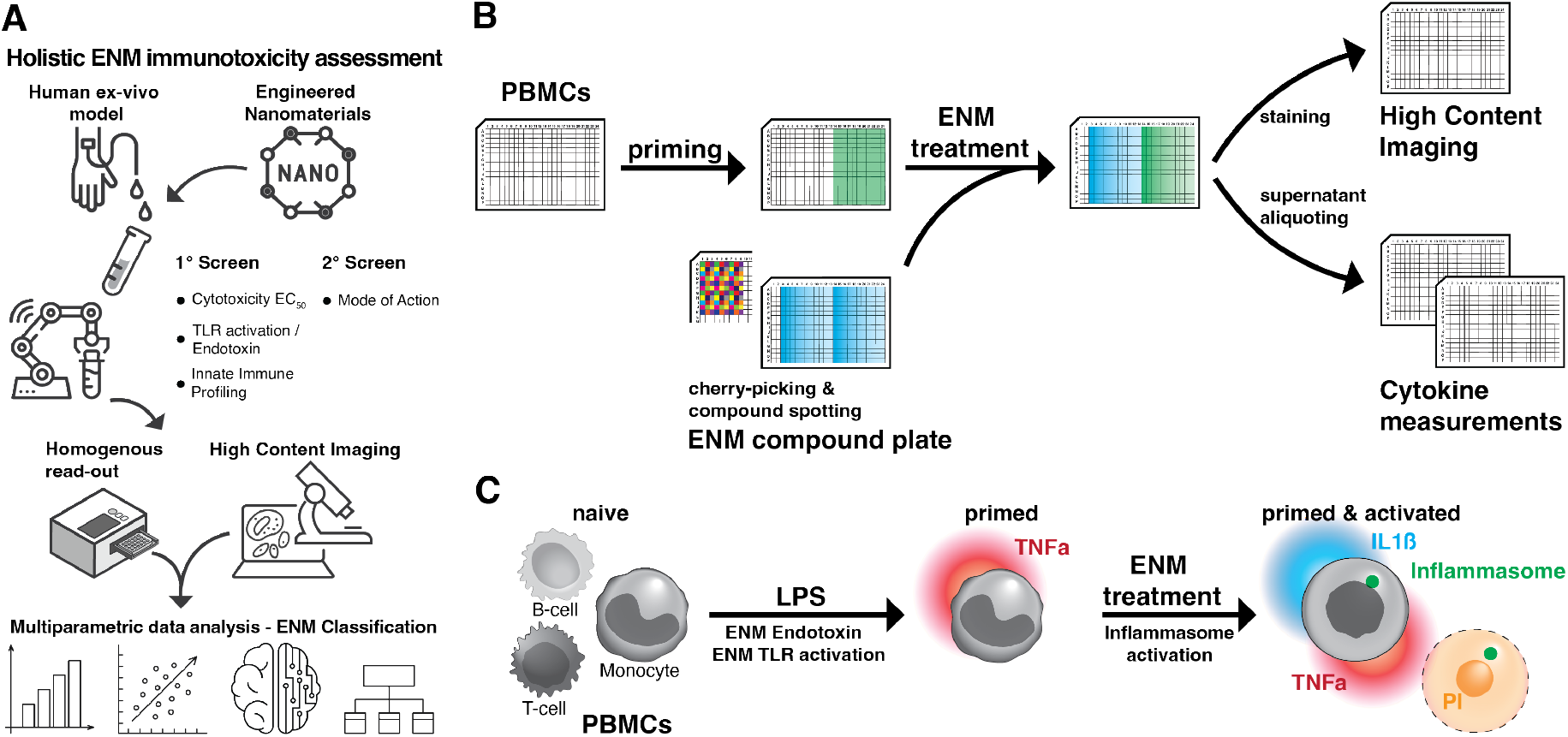
Schematic representation for (A) holistic ENM immunotoxicity assessment. With (B) practical workflows representing plate layout and (C) visualization of readouts on PBMCs.

**Figure 2:**
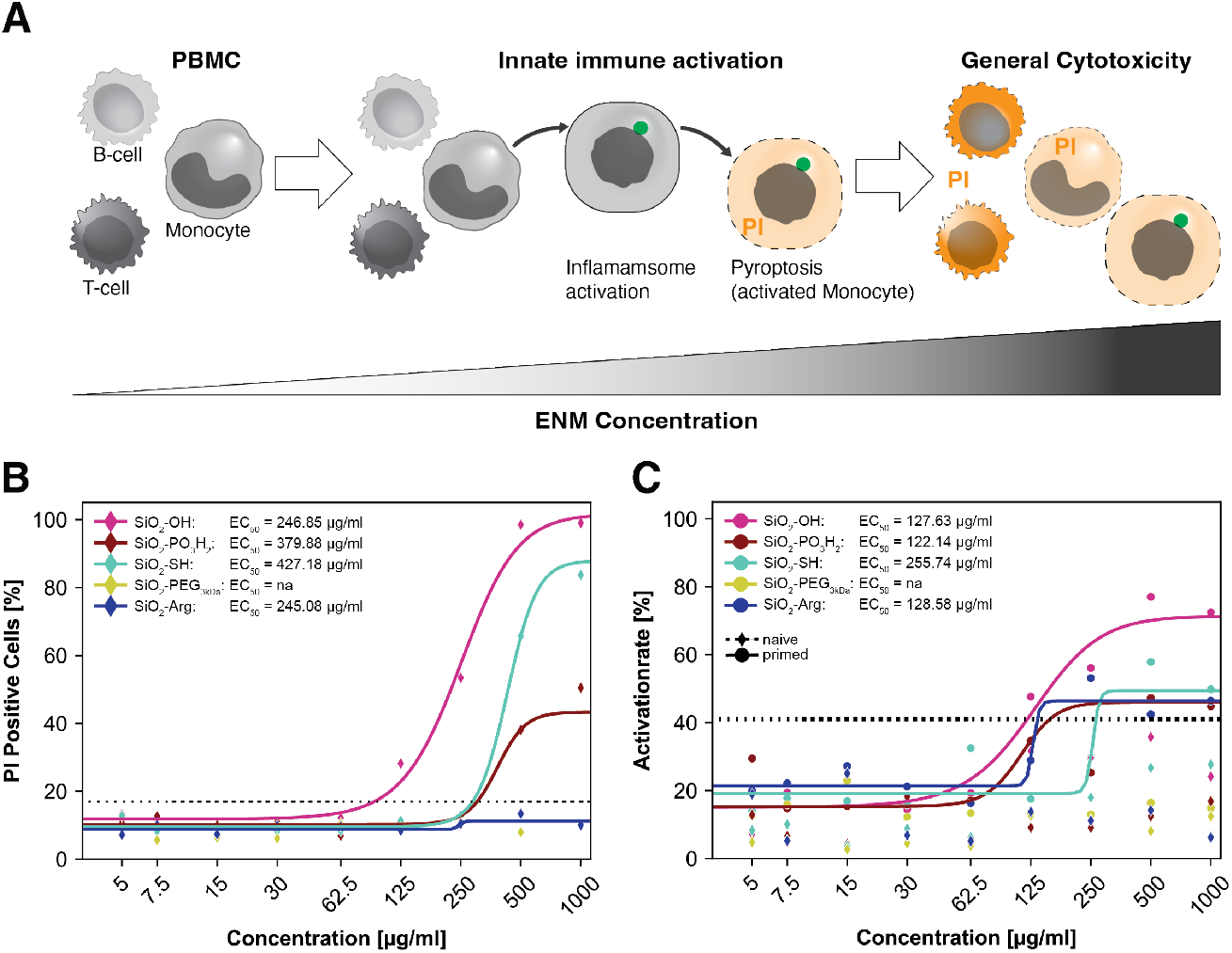
Cytotoxicity and innate immune activation. (A) by silica ENM with different surface functionalities. PBMCs response to Silica ENM (25 nm diameter) exposure (4 h treatment) was assessed in the naive state (2 h pre-incubation with medium) and in the primed state (2 h pre-incubation with LPS). Propidium iodide (PI, 5 µg/ml) was added for the last 30 min of the incubation time prior to fixation, after which cells were stained for inflammasome markers and analyzed by high-content imaging. Measurements (diamonds for the naive state and dots for the primed state) represent the mean of 16 image fields acquired. The lines show the result of a four-parametric logistic regression to determine EC_50_ values. The black dotted line shows the median + 3std as the activity threshold of non-ENM treated (B) naive control for PI-positive cells (17%) and (C) primed control for inflammasome activationrate (41%), n=48 wells with 16 images per well. (B) PI-positive cells are expressed as percentages relative to the total cell count. (C) Activationrate calculated as a percentage relative to CD14^+^ Monocytes.

### Cytotoxicity and innate immune response do (not) diverge

Next, we asked whether toxicity profiles of ENM-treated PBMCs in the naive (i.e., non-primed) state would translate to safe or hazardous innate immune activation. To investigate this, we looked into the cellular response of primed PBMCs to dose ranges of functionalized Silica ENM of 25 nm diameter. Modulation of the human innate immune response can be observed by combining image-based readouts from HCI as inflammasome activationrate (expressed relative to CD14 positive monocytes, see Figure S3) and PI-positive cells (Figure 2) with homogenous data of cytokine release (as shown later) at different states of the cells, e.g., naive and primed. When we compared the cellular response to ENM in terms of PI-positive cells in the two states (Figure S2A), we observed for some formulations (e.g., SiO_2_-Arg and SiO_2_-PO_3_H_2_) a general increase in PI-positive cells in the primed state for all ENM concentrations. Other formulations (-OH, - SH modifications) did not show a prominent difference between primed and naive states.

Silica ENM are known to activate the NLRP3 inflammasome ^[23]^. After NLRP3 inflammasome assembly and the consequent activation of caspase-I by its autocatalytic cleavage, cells enter pyroptosis, which is marked by permeable cell membranes ^[8]^. This means that using PI as a cytotoxicity marker will also label cells with activated inflammasome in pyroptosis, leading to the question of whether the observed differences in PI-positive cells of ENM-treated cells in the naive and primed state originate from those cells. Looking at the rate of inflammasome activation (Figure 2C) in the naive state (diamonds), no dose-dependent effects can be observed, suggesting that the tested formulations cannot trigger the TLR-4 receptor by themselves. In the primed state (dots), dose-dependent effects of inflammasome activation can be observed for -OH, -SH, and arginine-functionalized silica ENM, passing the threshold (mean plus three standard deviations of primed non-ENM control, 41%) with more than one data point. This data confirms the general capability of silica ENM to activate the inflammasome pathway. Notably, arginine-decorated ENM did not show a significant level of PI-positive cells, but significant inflammasome activation of primed PBMCs with EC_50_ of 129 µg/ml. This data shows the necessity to study immunotoxicity by combining features describing innate immune response (inflammasome activation with downstream IL1ß release) with a toxicity measure (PI-positive cells) over a dose range (rather than a single point) for each formulation in order to get insight for each individual formulation (Figure 3). In dotted lines thresholds of mean plus three standard deviations of the primed control are shown (activationrate: 41%, PI-positive cells: 21%, IL1ß release: 2232 pg/ml). Figure 3A shows that SiO_2_-OH ENM induces dose-dependent cytotoxic effects (EC_50, activationrate_=127 µg/ml) with correlated inflammasome activation. This activation exceeds thresholds at 125 µg/ml, reaching 99% PI-labelled cells at a dose of 500 µg/ml with more than 70% of monocytes mounting an inflammasome. At 125 µg/ml, 48% of monocytes show inflammasome activation with significant IL1ß release (6872 pg/ml), indicating functional inflammasome activation. With increasing ENM concentration, cytokine release of IL1ß (and TNFα, data not shown) drops. This drop is associated with the increasing and predominant toxicity effect of ENM ^[30]^. Arginine-functionalized silica ENM (Figure 3B) shows dose-dependent effects in inflammasome activationrate with an EC_50_ of 128.58 µg/ml and a maximum level of inflammasome activation of 53%. However, for IL1ß release and PI positive rate, only a moderate dose-dependent effect can be observed and only one data point at 500 µg/ml in PI positive rate and two data points (250, 500 µg/ml) surpass the defined threshold of three standard deviations above the primed control mean. Looking into the inflammasome response and the rate at which inflammasome-activated cells enter pyroptosis by inflammasome and PI-positive cells, we can see how the materials’ response can differ (Figure S2B). While for SiO_2_-OH of 25 nm size, both features highly correlate we can observe, that arginine-modified silica ENM mount a different response. Only about half of activated monocytes are also PI positive, while for SiO_2_-OH almost all activated cells are also PI positive (Figure S2B). This means SiO_2_-Arg activated monocytes can still be found with the impermeable cell membrane, while SiO_2_-OH activated monocytes are mostly found in the necrotic phase, indicating that we can assess the different states in the kinetic of inflammasome activation. When silica ENM are formulated with PEG (3kDa) (Figure 3C), no significant innate immune response could be detected. Measurements in all three parameters are kept within the threshold, and no dose-dependent effects can be observed.

**Figure 3:**
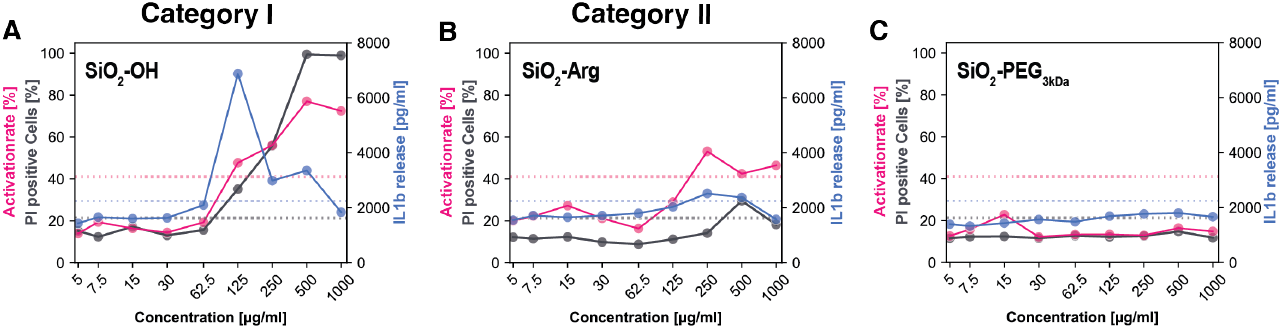
Innate immune activation profiling. Dose-dependent effects of 25 nm silica ENM on primed PBMCs. Individual graphs per formulation with a combination of activationrate [%] (pink, left y-axis), percentage PI-positive cells (gray, left y-axis), IL1ß release (blue, right y-axis). Thresholds of mean primed control + 3std in dotted lines (n=48, activationrate: 41%, PI-positive cells: 21%, IL1ß release: 2232 pg/ml). (A) SiO_2_-OH, 25 nm, (B) SiO_2_-Arg, 25 nm, (C) SiO_2_-PEG_3kDa_.

Two general categories of ENM can be identified by this method. Category I: ENM as SiO_2_-OH, -SH, and -phosphate (Figure S4) of 25 nm diameter which are toxic ENM with dose-dependent effects in all features, exceeding thresholds of the primed control with more than one data point. And category II: safe ENM that activate the innate immune system but do not induce exceeding cytotoxic effects, such as PEG-formulated silica ENM. While SiO_2_-Arg ENM fulfills the requirement for safe ENM, at a dose of 1000 µg/ml, cytokine release also drops, giving the indication of a predominant cytotoxic effect. At that concentration, therefore, this formulation must be considered toxic, category I.

### Particle size matters

Using the same method, we assess the size effect of SiO_2_-OH ENM-triggered inflammasome activation. Figure 4 depicts the innate immune activation pattern of silica ENM in diameters from 18 nm (Figure 4A) to 154 nm (Figure 4E). SiO_2_-OH ENM of 18-40 nm size (Figure 4A-C) can be classified as category I ENM, with PI-positive cells reaching 99% labeled cells from 500 µg/ml ENM concentration indicating general cytotoxicity. IL1ß release follows the same pattern for diameters between 18 nm and 68 nm. There is an increase in IL1ß release until it peaks and declines. The decline mostly correlates with predominant cytotoxicity. At 40 nm diameter, we can observe a lower maximum of IL1ß release in comparison to 25 nm diameter (drop from 6872 to 4961 pg/ml), with a peak shift from 125 µg/ml to 250 µg/ml ENM. This trend continues with increasing ENM diameter to 68 nm (Figure 4D), with IL1ß release maximum at 500 µg/ml ENM concentration, reaching 4313 pg/ml. For concentrations below 500 µg/ml, this ENM formulation shows functional inflammasome activation that does not exceed a toxic threshold, passing to toxic status from 500 µg/ml onward and reaching 63% PI-positive labeled cells at 1000 µg/ml. We, therefore, classify SiO_2_-OH ENM of 68 nm size as limited category II below 500 µg/ml. With increasing particle diameter to 154 nm, SiO_2_-OH ENM passes to category II, with only one data point passing the threshold for PI-labelled cells. Below 1000 µg/ml ENM concentration, no inflammasome activation can be detected. In summary, we can detect size-dependent effects of SiO_2_-OH ENM, detecting between 68 nm and 154 nm diameter the limit that ENM are safe for the human innate immune system.

**Figure 4:**
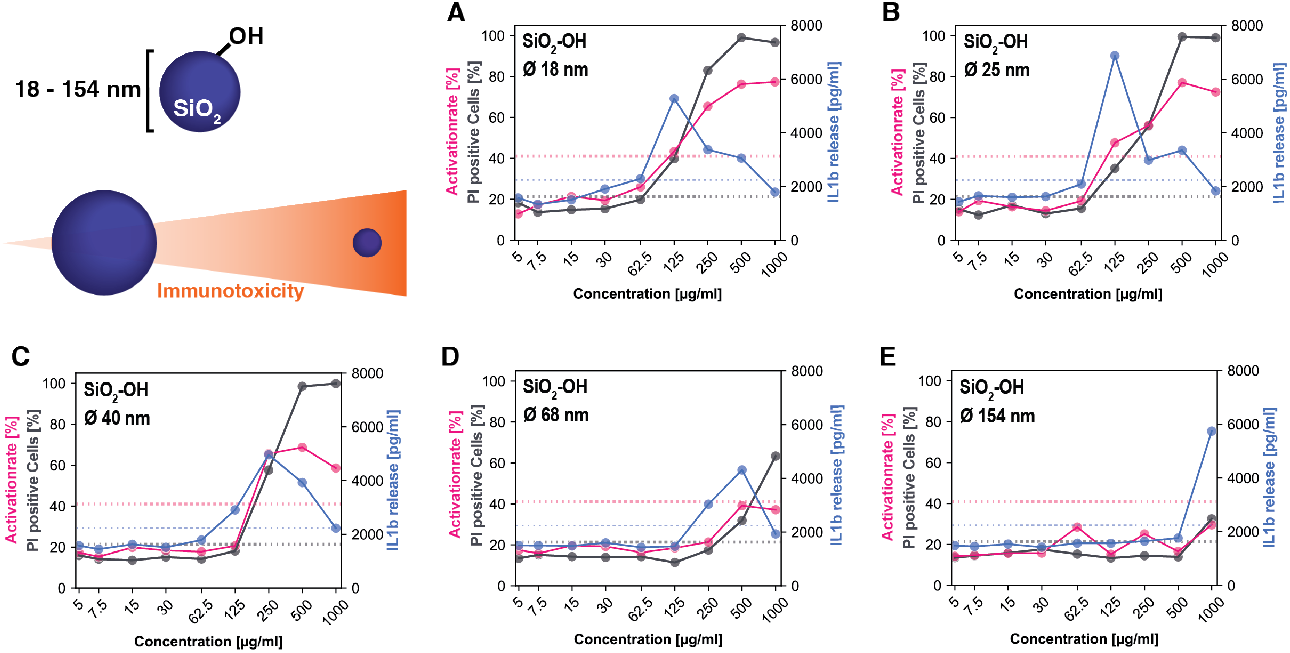
Size-dependent innate immune activation of SiO2-OH ENM. Dose-dependent effects of 25 nm silica ENM on primed PBMCs. Individual graphs per formulation with a combination of activationrate [%] (pink, left y-axis), percentage PI positive cells (gray, left y-axis), and IL1ß release (blue, right y-axis). Thresholds of mean primed control + 3std in dotted lines (n=48, activationrate: 41%, PI-positive cells: 21%, IL1ß release: 2232 pg/ml). (A-E) SiO_2_-OH, 18 – 154 nm diameter.

### Mode of action

To get mechanistic insights into ENM action in inflammasome activation, specific inhibitors were utilized (Figure 5). Inhibitors were added in a range of doses post LPS priming. 100 µg/ml SiO_2_-OH ENM (25 nm) were used for inflammasome activation for four hours. Subsequently, inflammasome activation, cytokine expression, and release were analyzed from cell-based high-content imaging data and homogenous time-resolved fluorescence (HTRF) data. Using MCC950, a small molecule inhibitor for the NLRP3 inflammasome, 75% inhibition (EC_50_ = 0.28 µM) of inflammasome activation was achieved with 95% inhibition of IL1ß release at EC_50_ of 0.18 µM (Figure 5A). NLRP3 inflammasome assembly triggers caspase-I cleavage, which is required for the maturation of pro-IL1ß and release of IL1ß to the cytosol. Using the caspase-I inhibitor VX-765, we could not observe dose-dependent changes in inflammasome activation (Figure 5B) but complete inhibition of IL1ß release from 20 µM inhibitor concentration with EC_50_ of 2.1 µM. This data indicates that SiO_2_-OH ENM triggers the NLRP3 inflammasome activation with caspase-I dependent release of mature IL1ß. Next, we asked which way SiO_2_-OH ENM activates the NLRP3 inflammasome, therefore, we assessed the role of the P2X7 receptor and cathepsin B. If the ENM cause cellular stress and release of ATP, the P2X7 receptor, sensing extracellular ATP, would be actively involved in the signaling cascade of SiO_2_-OH ENM-mediated inflammasome activation. However, the addition of the P2X7 receptor antagonist A-438097 in a dose range from 0.29 to 45.7 µM, does not significantly influence inflammasome activation nor IL1ß positive cells or IL1ß release (Figure 5C). This indicates that (i) the P2X7 receptor is not involved in SiO_2_-OH ENM-induced inflammasome activation, and indirectly, that (ii) 100 µg/ml SiO_2_-OH ENM do not trigger stress-induced ATP secretion that in turn activates P2X7 receptor. However, when inhibiting inflammasome activation through cathepsin B, which is released from ruptured lysosomes, we can observe a dose-dependent reduction of inflammasome activation (with EC_50_ at 3.9 µM and 70% maximum inhibition) and complete inhibition of IL1ß release from 40 µM inhibitor concentration (EC_50_ = 3.38 µM), correlating with an increase in intracellular IL1ß (Figure 5D). This data suggests that SiO_2_-OH ENM are activating the inflammasome through the lysosomal pathway.

**Figure 5:**
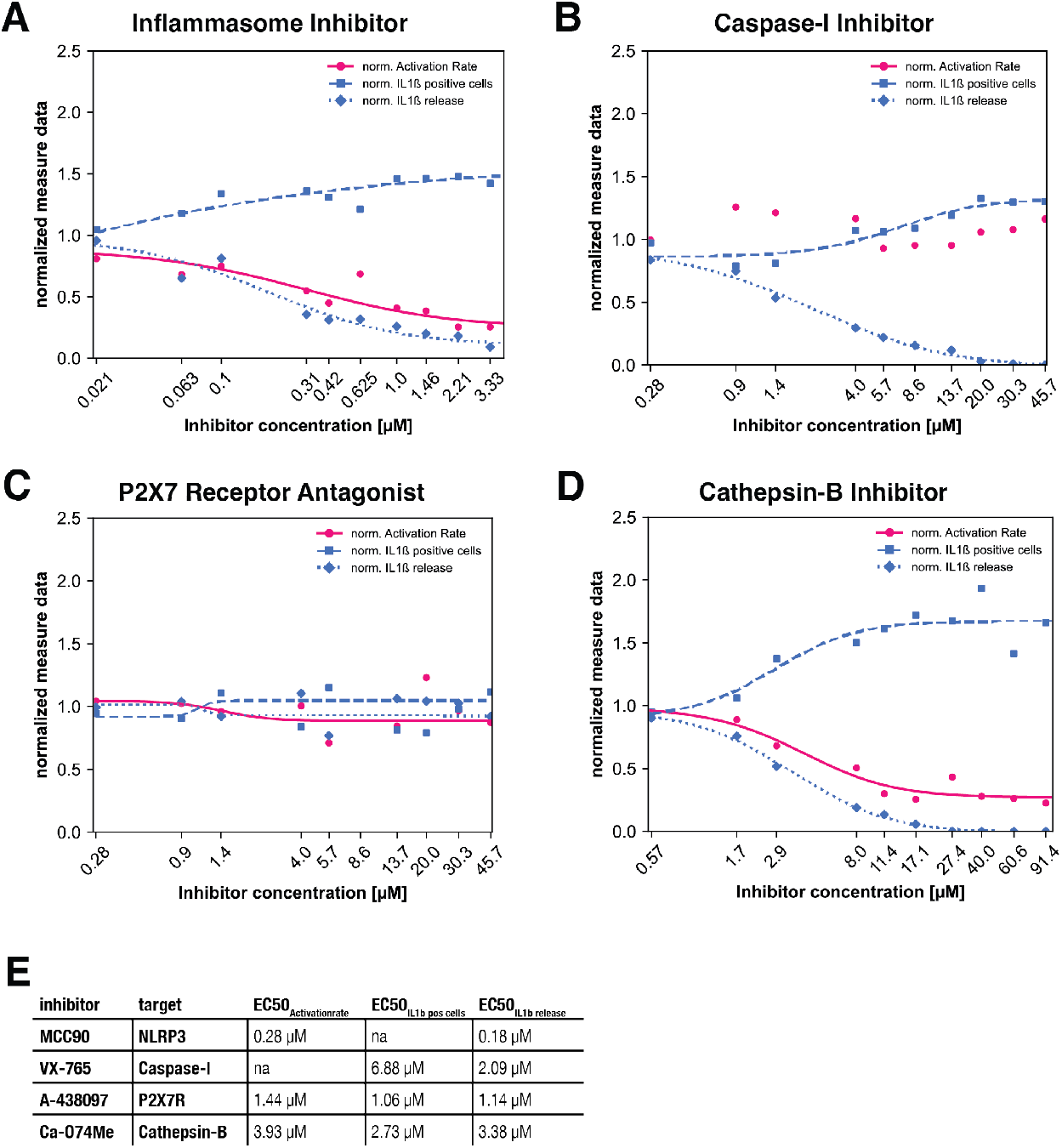
Mechanistic insights into Inflammasome activation through SiO2-NPs. PBMCs were primed for 2 h with LPS after which inhibitors at different doses were added for one hour. SiO2-OH ENM (25 nm diameter) at 100 µg/ml were added for innate immune activation. (A) MCC90 as an inflammasome inhibitor, (B) VX-765 as a caspase-1 inhibitor, (C) A-438097, a P2X7 receptor antagonist, and (D) Ca-074Me, a cathepsin-B inhibitor were used in this assay. Quantification of activationrate (pink, solid line), IL1ß positive cells (blue, dashed line) from HCI, and IL1ß release (blue, dotted line) from HTRF are shown as points. Results of four parametric log-logistic regression to calculate EC_50_ values (E) are drawn as curves.

### Endotoxin contamination of ENM

As it has been shown by others, endotoxin contamination on ENM are underestimated ^[19, 20]^. Those contaminations originate mostly in the manufacturing process. To detect endotoxin contamination on ENM, different assays can be performed. Most prominent LAL and Gel Clot endotoxin assays are performed. They are spectrophotometric methods and underlay the experimental bias through light absorption and diffraction through ENM. Components of endotoxins bind to TLR-4, triggering intracellular TNFα and pro-IL1ß expression in monocytes with release of TNFα, reflecting the first stimulus in the pathway to inflammasome activation. Using LPS as an endotoxin model on a fixed dose of SiO_2_-OH ENM (25 nm, 100 µg/ml), we show that TNFα release (Figure S5A) responds to LPS in a dose-dependent manner. Using four parametric log-logistic regression, the half maximal effective concentration of LPS was calculated for both features. The Limit of Detection (LOD) and Limit of Quantification (LOQ) were obtained using the blank method, where LOD is defined as mean plus three standard deviations and LOQ as mean plus ten standard deviations ^[39]^. Therefore, the mean activity and standard deviation of the blank (no LPS, no ENM) were calculated to obtain LOD and LOQ activity levels. LPS concentrations of LOD and LOQ were calculated at the corresponding activity levels by the curve fit parameters defined by the four parametric log-logistic regression, as seen in Figure S5B. The data show that LPS concentrations on SiO_2_-OH ENM with a limit lower than 0.05 ng/ml (corresponding to 0.05 endotoxin units, EU, per ml) can be detected in this assay, which is competitive to other methods ^[40]^, indicating that, using this assay, endotoxin contamination on ENM can be detected and discriminated from an ENM-mediated effect on inflammasome activation. To identify endotoxin-contaminated ENM, TNFα release in the naive state must be above the threshold of mean plus three standard deviations of the naive control. As we show in Figure 6A, TNFα release in the naive state for all formulations and concentrations was measured below the threshold, and no dose-dependent effects could be determined. No endotoxin contamination was detected on the tested materials.

**Figure 6:**
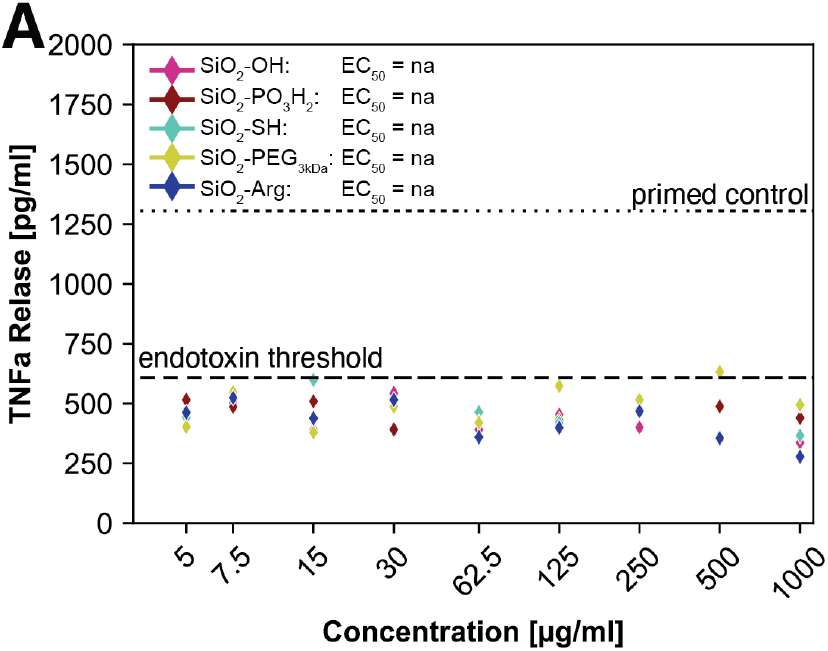
Orthogonal Screen of Endotoxin contamination on ENM. (A) TNFα release [pg/ml] measured in the supernatant of ENM treated PBMCs in the naive state (2 h pre-incubation with medium with following treatment of ENM at different doses). Dashed lines denoted as endotoxin threshold shows mean+3std of the naive control, considered as threshold of endotoxin contamination. Dotted lines depict the median of the primed control (n=48 wells with 16 image fields per well).

## 3. Discussion

ENM of different materials were shown to trigger an inflammatory response, leading to or aggravating inflammatory conditions in humans ^[5, 9, 11]^. In particular, metal oxides such as silica ^[18, 21, 23-26]^, TiO_2_^[24, 27]^, and CeO_2_^[28]^, among other ENM ^[29-32]^, were shown to activate the innate immune system via NLRP3 inflammasome. The broad activation of the innate immune system may directly result in adverse inflammatory reactions, but it was also shown that inflammasome activation can modulate disease progression. In the case of Alzheimer’s disease, activation of the NLRP3 inflammasome pathway in microglial cells leads to the release of inflammasomes to the extracellular space ^[41]^. These inflammasomes can bind to amyloid-beta monomers, increasing their probability of aggregation and, by that further to the progression of Alzheimer’s disease ^[42]^. This clearly shows the need to test inflammasome activation as an immunosafety endpoint.

Small molecules are in contrast to ENM well regulated. However, regulatory guidance for immunotoxicity testing is poorly available for small molecules ^[3]^. Nevertheless, it is currently the only guidance documentation used for ENM. In our view, this is insufficient as the special physicochemical characteristics of ENM along with different interactions at the nano-bio-interface define ENM as a unique material. Thus, it is necessary to develop standard test methods in relevant human-based models. To keep pace with the rapid pace of new ENM development ^[6]^, high-throughput methods must be developed for scalable ENM toxicity testing that can contribute to the development of new test guidelines ^[3-4, 7]^.

Our assay shows a comprehensive *ex vivo* high-throughput screening system for ENM allowing (i) high sample throughput, (ii) assessment of dose-response relationships, (iii) multiparametric readout for cytotoxicity, and innate immunity at an early time point in a combinatorial method to avoid NP-induced bias by (iv) orthogonal endotoxin screen.

By testing various surface-modified silica ENM of the same size (25 nm), we observed differences in cytotoxicity in the naive state (Figure 2B) and inflammasome activation in the primed state (Figure 2C). From our results, we distinguished two categories of ENM: Category I ENM, inducing general cytotoxicity and inflammasome activation, and Category II ENM, which activate the innate immune system but do not show general cytotoxic effects. We verified NLRP3-specific inflammasome activation (Figure 5A, B) that is triggered through the lysosomal pathway with the release of cathepsin B (Figure 5D), which is most prominent for particulate material ^[10]^. This means that the inflammatory reaction is induced by NLRP3 inflammasome activation and not by the release of DAMPs like ATP. Unmodified (SiO_2_-OH), thiol-modified, and phosphate-modified silica ENM fall under the Category I profile, yet with varying cytotoxicity levels reached (SiO_2_-OH >, -SH > -PO_3_H_2_, Figure 2). Although SiO_2_-Arg did not show cytotoxicity, dose-dependent effects for inflammasome activation were measured up to a significant level. This immunomodulatory effect can contribute to disease progression and classifies SiO_2_-Arg as a category II ENM. In a traditional assay, only Category I ENM would be identified as harmful, but Category II ENM may also be identified as harmful if tested in a sufficiently high concentration (e.g., 1000 µg/ml). This demonstrates the need to assess immunotoxicity in different states and from various perspectives.

When comparing different sizes of ENM, smaller ENM of the same mass concentration have (i) a higher particle concentration, yielding different amounts of internalized substances ^[43]^, and (ii) a relatively higher reactive surface area, leading, e.g., to higher ROS for smaller ENM in comparison to larger ones *in vitro* and *in vivo* ^[44]^. Inflammasome activation by polystyrene ^[31]^, TiO_2_ ^[24]^, and silver ^[30]^ ENM has been shown to be size-dependent. Small (below 20 nm) silica ENM were shown to induce higher levels of IL1ß compared to larger (100-200 nm diameter) silica ENM on human PBMCs ^[25, 45]^. When assessing the immunotoxicity profile of silica ENM between 18 and 154 nm, we could observe the same trend. There is a visible drop in immunotoxicity from 25 nm to 40 nm, leading to a safe profile at 154 nm diameter (Figure 4), where cytotoxicity and innate immune activation are kept below the threshold until a concentration of 500 µg/ml.

However, our data show that at the given size of 25 nm SiO_2_-OH particle elicits a toxic effect that reaches about 56% at 250 µg/ml. In contrast, the SiO_2_-Arg decorate particles do not trigger significant toxicity at 250 µg/ml (15%, Figure 3). Even at the maximum dose of 1000 µg/ml, the SiO_2_-Arg ENM does show a safe toxic profile. This data clearly show that size alone is not sufficient to predict toxicity, but rather the combination of size with surface properties is the cause of cytotoxicity. This has an impact on the de-risking of ENM as surface functionalization can abolish the toxic effects of ENM of a certain size and it needs to be considered in the safe-by-design criteria.

The data shows that using our platform, we can discriminate between cytotoxicity and inflammasome activation. In summary, we show that the immunotoxicity effects of ENM are composed of (i) innate immune activation and (ii) general cytotoxic effects. Therefore, it is necessary to tune the assay platform to perform multiparametric analysis in order to understand the immunotoxicity of ENM.

Our high-throughput screening concept is based on assessing the human innate immune response at an early stage. Typical experimental protocols with THP1 cells involve long experimental protocols (18-24 h), which are necessary to induce the primed state of THP1 cells. Long-term treatment of monocytes/macrophages (>6h) increases the probability of non-canonical inflammasome activation, meaning mixed mechanisms (canonical and non-canonical) of inflammasome activation contributing to IL1ß release ^[46, 47]^. Innate immunity reactions are by their nature set to be fast reactions that happen in seconds/minutes from the triggering of the event. This is the physiological response of innate immunity. True is that in chronic conditions innate immunity can be dysregulated and be continuously activated. However, it is very challenging to model this phenomenon *ex-vivo* as the machinery for the continuous production of new PBMC is missing. Hence, if a chronic exposure setting (24 h cell-material contact) on PBMCs is chosen, innate immunity readouts are masked and are not the correct endpoint. Using the experimental protocol reported here, early innate immunity response after four hours (or less) of cell material contact can be read, enabling the assessment of inflammasome activation kinetics and the fate-activated monocytes undergo (inflammasome activation - pyroptosis - necrosis, Figure S2B). Our screening assay is miniaturized to 384-well format, fully automated and scalable to test a plethora of ENMs. The assay is performed in dose-response format, which is important to calculate threshold limit values of hazardous ENM by quantitative dose-response relationships. Thus, the here provided method gives risk assessment data. For some ENM, we show that lower doses are safe, whereas higher doses cause severe cytotoxicity. If such ENM are considered for application in a pharmacological context, the obtained data can already indicate potential side effects through local accumulation. By four parametric log-logistic regression on the dose-response data, we can identify dose-dependent effects and calculate EC_50_ parameters. Our assay allows the obtaining of quantitative time-response relationships, assessing the development of the innate immune reaction through measurements at different states. Assay performance measures based on Z’-score, intra-, and inter-plate variance (Table 1, Figure S1) show the high quality and scalability of our screening assay. Measurements at different states and the combination of different readouts (homogenous and single cell-based high-content imaging data) allow us to obtain an integrated insight into ENM-caused immunotoxicity effects by conducting a single experiment. With multiparametric analysis of this high-dimensional data, the mode of action of a given ENM can be determined and classified, as we show exemplary for unmodified silica ENM (Figure 5). This serves as a harmonized and holistic safety assessment of immunotoxicity effects caused by ENM. Additionally, this assay can be used in ENM/drug development processes in a safe-by-design and production-to-product approach.

### Limitation & Outlook

Our experimental design was based on silica-based ENM. Silica ENM are largely used in several sectors like construction, cosmetics, and medicine and our rationale was to use them as a representative class of ENM for this study. However, we are aware that different classes of ENM exist and that they also need to be tested using this approach.

The screening assay described in this work is based on primary human cells with underlying biological variation. As we said we take into consideration this aspect by referring to the sample tested with our database. However, we believe that it is of pivotal importance to use human primary cells in the assessment of ENM safety to correctly predict their effects on human health. An example of a potentially dangerous underestimation of the negative effect of ENM was reported by Tahtinen et al. In this study, the authors showed the severity of host-dependent systemic inflammatory responses by liposome-based mRNA nanomedicines ^[37]^. Inbred laboratory mice tolerate 1000-fold higher doses of vaccine in comparison to humans, who react already at lower doses of RNA vaccine with adverse side effects, e.g., fever and chills, through the systemic increase of pro-inflammatory cytokines ^[37, 48]^. Using human-based systems, adverse events in pre-and early clinical phases could be avoided. The inherent biological donor variation is not a throwback for the assay but an essential part. It is known that the composition of cell types in the blood can differ between women and men and varies with age ^[49, 50]^. It is also known that immune responses, e.g., to vaccination, differ for obese and non-obese individuals, meaning it is dependent on the body mass index ^[51]^. The use of primary cells in this screening system allows us to exactly understand the influence of a compound, for instance, on a group of subjects at risk such as obese individuals. The strategy, therefore, is rather to select donors by parameters of interest and to choose then a set of representative donors from our donor database (data not shown) to evaluate innate immune effects for a distinct stratified group.

In this way, the here-described integrated immunotoxicity assay for ENM can be used (i) for safety assessment and (ii) in the development of safe-by-design ENM.

In conclusion, we developed an assay capable to detect the cytotoxicity and immunotoxicity of ENM in a specific manner. Our assay is based on human PBMC and therefore is highly relevant to the ENM safety assessment. As the assay here proposed is fully automated and miniaturized our assay is amenable for large screening campaigns and is satisfying the requirement of the majority of the regulatory agencies. In addition, our assay is capable to give not only information about the immunotoxicity elicited by ENM but also their cytotoxicity and data about endotoxin contamination. Additionally, we showed that our assay gives the possibility to the investigator to understand and deconvolve the mechanism of action of the ENM effect on the PBMC. This is an added value as the data obtained can be used to troubleshoot the ENM or in case of biomedical use, the ENM can be tuned in the desired direction. As our assay can be combined with other read-outs, like cytokines detection, proteomics, transcriptomics, and other omics data we are enabling the possibility to have a deep-phenotype approach that can be extremely useful in the development of biomedical ENM. In essence, we believe that our assay represents a significant advancement in the field of nanotoxicology and biosafety assessment in a systematic and screenable format. The assay presented here stands for the future development of a novel class of cell-based assay dedicated to the test of ENM.

### Experimental Section

#### PBMCs isolation

Peripheral Blood Mononuclear Cells were isolated from buffy coats obtained from the German Red Cross by density gradient centrifugation. In brief, buffy coats at room temperature (RT) were diluted 0.75x with PBS and gently homogenized. Next, falcons containing Pancoll (PAN Biotech) were overlayed with the diluted buffy coats and centrifuged for 20 min (700 g, RT) without brakes. Mononuclear cells were collected from the plasma/Pancoll interface and washed several times with PBS for purification. Cell number and viability was determined using Vi-CELL XR (Beckman). Cells were then aliquoted in 50 million portions and stored in a liquid nitrogen vapor phase storage system until further usage.

#### ENM inflammasome activation assay

PBMCs of one donor were selected and reconstituted in RPMI medium (Biochrom) containing L-Alanyl-Glutamine (200 mM, Biochrom), HEPES (1 M, LifeTechnologies), Ciprofloxacin (200 mg/ml, Kabi) and Hi-FBS (10%, Sigma) after thawing. A cell suspension of 1.5×10^6^ cells/ml was prepared and added to each well of a 96 deep-well plate (Thermo-Nunc, 1.0 ml). For cell seeding, the Biomek i7 Automated Workstation was used to dispense 40 µl cell suspension (60.000 cells/well) into each well of a black 384-well microplate with transparent bottom (Perkin Elmer) suitable for high-content imaging. Next 20 µl LPS (20 ng/ml InvivoGen) or medium was added to the wells and incubated for 2 h at 37°C and 5% CO_2_. For all steps (priming, activation, staining), polypropylene deep-well source plates were prepared with reagent or medium accordingly to the plate layout. Pipetting steps were all conducted using an i5 automated workstation with a 384 pipetting head and pipetting tips of 25 or 50 µl from Beckman. The compound plate (PorVair, low volume) with different doses of ENM (SiO_2_ ENM of different sizes and surface modification, obtained by SILVACX) and solvent control was prepared by transferring respective volumes using an acoustic liquid handler (Echo 525, Beckman). Therefore, source plates (Labcyte-PP-0200) were filled with 50 µl of each ENM formulation (20 mg/ml) and water as solvent control. Samples are then diluted with PBS/HEPES (25 mM) up to 10 µl on a Multidrop (Thermo). 5 min prior to compound addition, samples are placed on an orbital shaker for 5 min. After 2 h of incubation with LPS, ENM and solvent controls (in different doses) were added and incubation was continued for more than 3.5 hours when 10 µl of Hoechst/propidium iodide (H33342: 2 µg/ml, PI: 5 µg/ml, Sigma-Aldrich) staining and ATP (2 mM final concentration, Sigma-Aldrich) activator control was added. After 30 min of incubation, supernatants were removed and aliquoted for cytokine measurements. Cells were washed four times with PBS containing 1% FBS and cells were fixated with PFA (4%, methanol-free, Thermo) subsequently for 15 min. After removal of PFA, PBS containing 1% FBS was added and the plate was sealed and stored at 4°C.

#### Inhibitor study

To understand the pathways involved in inflammasome activation, a similar assay as the ENM inflammasome activation assay was performed on the same technological infrastructure. Briefly, 60.000 human PBMCs per well were seeded to each well of a black 384-well microplate with a transparent bottom (Perkin Greiner) suitable for high-content imaging. Subsequently, 20 µl LPS (final concentration: 20 ng/ml InvivoGen) or medium was added to the wells and incubated for 2 h at 37°C and 5% CO_2_. A compound plate with different stock solutions of inhibitors - MCC950 (20 mM, Cayman Chemical), TAK-242 (20 mM, Cayman Chemical), A-438097 (20 mM, Selleckchem), VX-765 (10 mM, Selleckchem) and Ca-074ME (40 mM, Enzo) – in dimethylsulfoxide (DMSO, Merck) was prepared by using an acoustic liquid handler (Echo 525, Beckman). 10 µl of inhibitor compound (or solvent control) was added to the assay. After one hour of incubation, SiO_2_-OH ENM (25 nm, Silvacx) in a final concentration of 100 µg/ml were added to the cells as activating agent for 4 hours. No-inhibitor, non-ENM treated control conditions received ATP (final concentration 2 mM, Sigma) as activating agent for 30 min. After removing and aliquoting cell supernatants for cytokine measurements, cells were washed with PBS containing 1% FBS and fixated with PFA (4%, methanol-free, Thermo) subsequently for 15 min. After the removal of PFA, PBS containing 1% FBS was added and the plate was sealed and stored at 4°C.

#### Immunocytochemistry

To prepare cells for high-content imaging, immunocytochemistry (ICC) staining was performed. Therefore, blocking and permeabilization buffer containing 1% Bovine Serum Albumin (Sigma) and 0.5% Triton-X-100 (Sigma) in PBS (DPBS w/o Ca^2+^/Mg^2+^, Gibco) was added to the cells for 20 min at RT. Next, the ICC staining solution was prepared by adding the respective antibodies to the blocking and permeabilization buffer containing 1 µg/ml Hoechst33342 (Sigma). In the case of the ENM inflammasome activation assay, Anti-Pycard-Dy488 (1:500, LSBio) was used. In the case of the Inhibitor study, Anti-IL1-beta-AF594 (1:100, R&D Systems) was used additionally. After gentle homogenization, ICC staining solution was added to the cells and incubated for 1 hour in the dark at RT. Cells were then washed with PBS containing 1% FBS, and the plate was sealed and stored at 4°C until image acquisition.

#### High-content imaging

Cells were imaged on an automated confocal spinning-disk fluorescence microscope CV6000 and CV8000 (Yokogawa) using a 40x water objective. 16 image fields per well were recorded and stored with maximum intensity projection of a z-stack with 3 layers.

#### Image analysis

Quantifications from High-Content Microscopy were performed using Yokogawa CV7000 Software. Therefore, at first nucleus detection was performed. Based on ENM-free controls in naive and primed conditions, a threshold for PI-positive and PI-negative cells was defined. In the case of IL1ß staining, a threshold for IL1ß positive cells was defined based on the ENM-free controls in naive and primed conditions. In order to obtain an artificial cell mask, the nucleus mask was dilated. Inflammasome spots were detected by object detection. Inflammasome-positive cells were identified, by dividing cells into populations containing (or not) inflammasome objects inside the cell mask. Rates of positive cells were calculated through the percentage of positive cells from the total number of cells per well.

#### Cytokine quantification

To quantify the cytokine concentration in the supernatants of the cells, HTRF measurements of IL1ß and TNFα were conducted according to the manufacturer’s instructions.

#### Dose-response analysis

To obtain EC_50_ values from dose-response data, four parametric log-logistic regressions^[52]^ were performed using the formula:

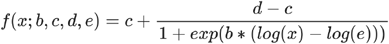

With parameters b: Hill slope, c: minimum response, d: maximum response and e: EC_50_. Two requirements needed to be met in order to be considered well-fit. (i) The EC_50_ is within a limit of the tested dose range (between 7.5 and 500 µg/ml), and (ii) the residuals < 400. (Note: non-normalized data was used to be fit which lets the residuals seem high in here.)

## Acknowledgments

The authors acknowledge the team of the CRFS, Gisela Schmidt and Sven Fengler for the critical revision of the manuscript.

