## Supporting Information for "An ex vivo human model for safety assessment of immunotoxicity of engineered nanomaterials"

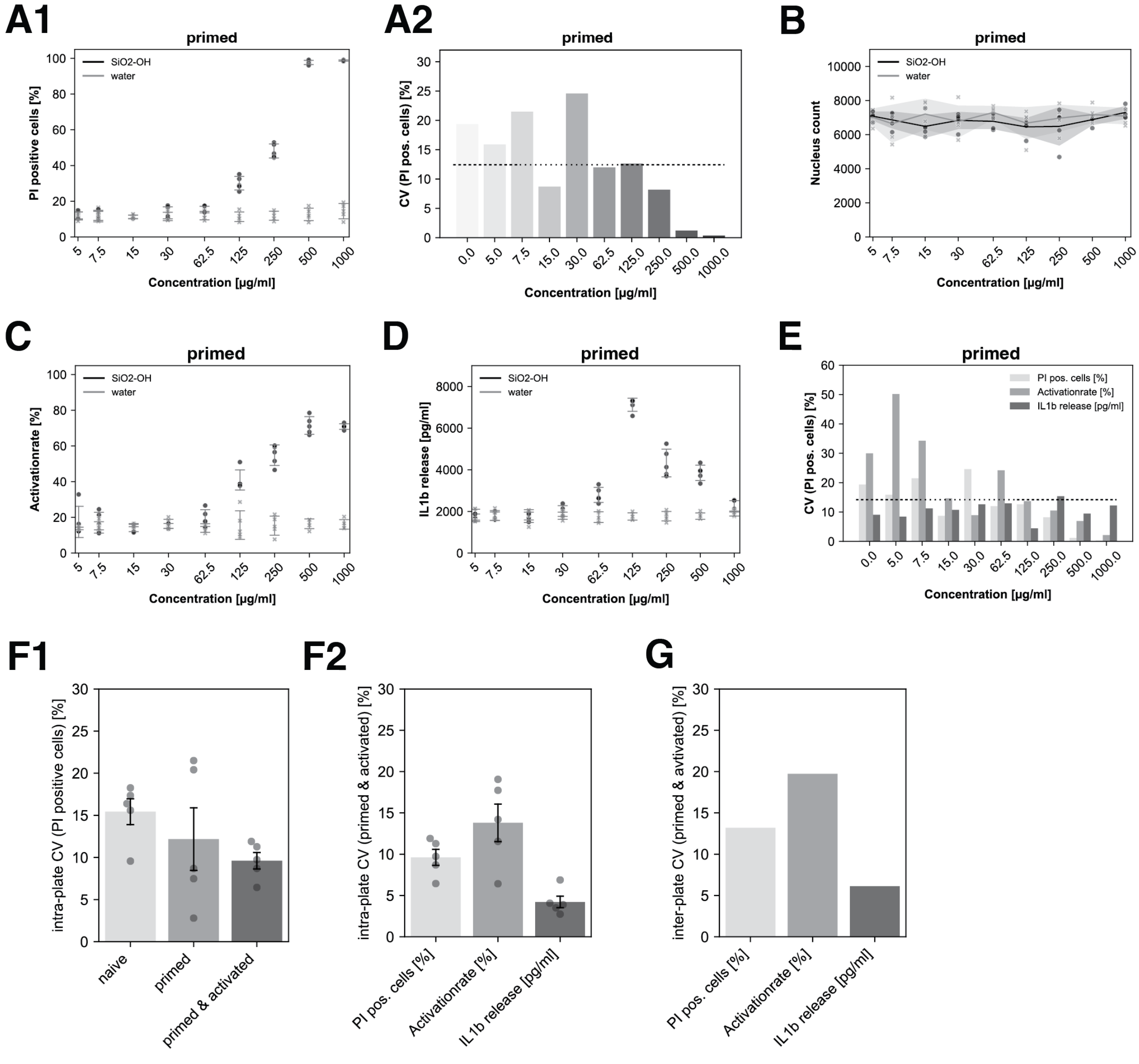


***Figure S1:*** **Assay performance measures.** Each screening plate contains a section of reference ENM (A-F) and non-ENM (G-H) controls. As a measure for assay robustness, the coefficient of variation, Z'-score, and assay window are calculated. For reference, ENM ($SiO_2$-OH, 25 nm) and solvent control similar dose ranges as for the tested ENM are prepared. (A-E) show data of different measures, collected on water solvent control and reference ENM for the primed state. Error bars represent the standard deviation of n=5 plates. (A2) inter-plate variance of PI positive cells in the primed state of reference ENM treated cells for each concentration. The black dotted line shows the mean CV over all concentrations (12%). (E) inter-plate variance of measured parameters by reference ENM concentration in the primed state. Mean CV (14%) over the three features and all concentrations on five plates as black dotted line. Inter-plate (F) and intra-plate (G) variation for non-ENM controls. Intra-plate variance of (F1) PI positive cells in naive, primed and primed & ATP activated state. (F2) Intra-plate (n=8 wells on n=5 plates, data shown as mean ± SEM on error bar) and (G) inter-plate variance (n=40 wells of n=5 plates) of IL1ß release, PI positive cells and activationrate in the primed & ATP activated state.

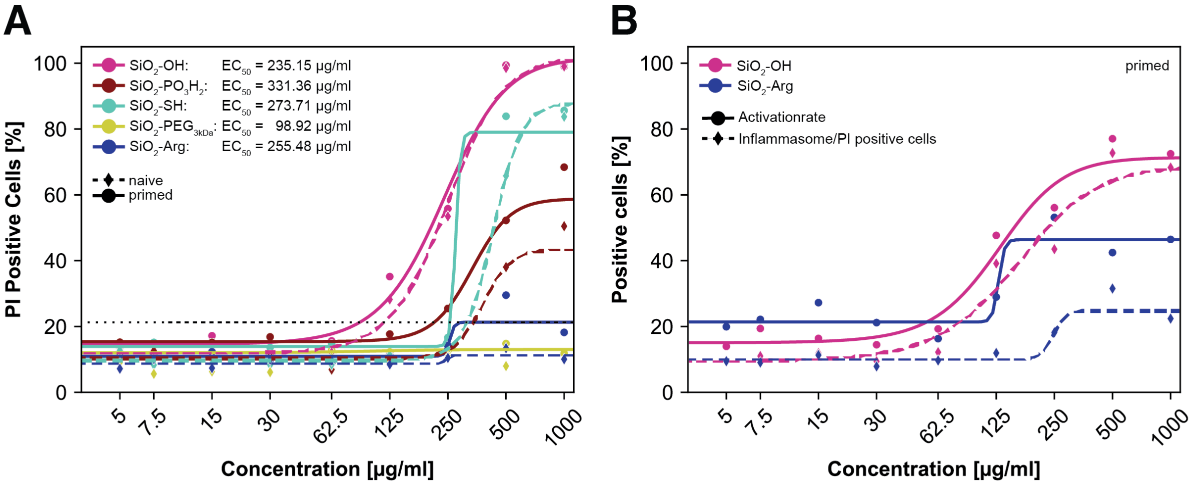


***Figure S2:*** **Cytotoxicity of Silica ENM with different surface functionalities in naive and primed state.** PBMCs response to Silica ENM (25 nm diameter) exposure (4 h treatment) was assessed in the naive state (2 h pre-incubation with medium) or in the primed state (2 h pre-incubation with LPS). Propidium iodide (PI, 5 µg/ml) was added for the last 30 min of the incubation time prior to fixation and analysis by high-content imaging. Measurements (diamonds for the naive state and dots for the primed state) represent the mean of 16 image fields acquired. The lines show the result of a four parametric logistic regression (dotted for the naive state and solid for the primed state) to determine $EC_{50}$ values. The black dotted line shows the median of primed controls + 3 std as activity threshold (n=48 wells with 16 images per well). (A) PI positive cells are expressed as percentage relative to the total cell count. (B) Inflammasome activationrate (dots, solid line) and Inflammasome/PI positive cells (diamonds, dashed line) in the primed state for $SiO_2$-OH and $SiO_2$-Arg ENM with 25 nm diameter.

**A**

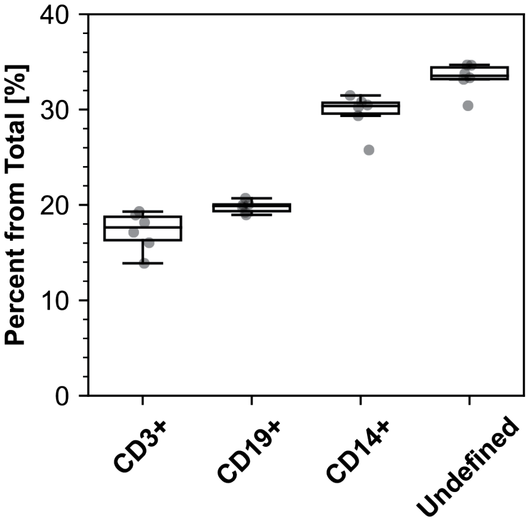


**B**

| Percentage of Cell Type | mean | std | cv | 25% | 50% | 75% |
|---|---|---|---|---|---|---|
| **CD3+ Cells [%]** | 17.2 | 2.0 | 11.8 | 16.3 | 17.6 | 18.8 |
| **CD19+ Cells [%]** | 19.8 | 0.6 | 3.2 | 19.4 | 19.9 | 20.1 |
| **CD14+ Cells [%]** | 29.7 | 2.0 | 6.9 | 29.6 | 30.4 | 30.7 |
| **Undefined Cells [%]** | 33.3 | 1.6 | 4.7 | 33.2 | 33.6 | 34.4 |

***Figure S3:*** (A) Cell type fractions of Monocytes (CD14+), B-cells (CD19+), T-cells (CD3+) and an undefined population, quantified from ICC. (B) Quantification of the same cell types by flow cytometry.

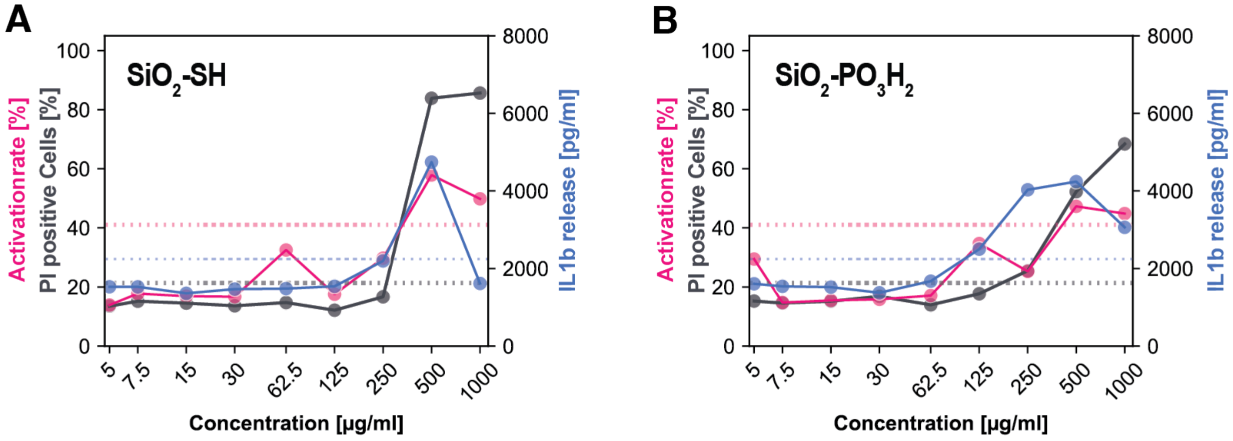


***Figure S4:*** **Surface modification of Silica ENM with 25 nm diameter affects innate immune response.** Dose-dependent effects of 25 nm silica ENM on primed PBMCs. Individual graphs per formulation with a combination of activationrate [%] (pink, left y-axis), percentage PI-positive cells (gray, left y-axis), IL1ß release (blue, right y-axis). Thresholds of mean primed control + 3std in dotted lines (n=48, activationrate: 41 %, PI-positive cells: 21 %, IL1ß release 2232 pg/ml). (A) $SiO_2$-SH, 25 nm, (B) $SiO_2$-$PO_3H_2$, 25 nm.

**A**

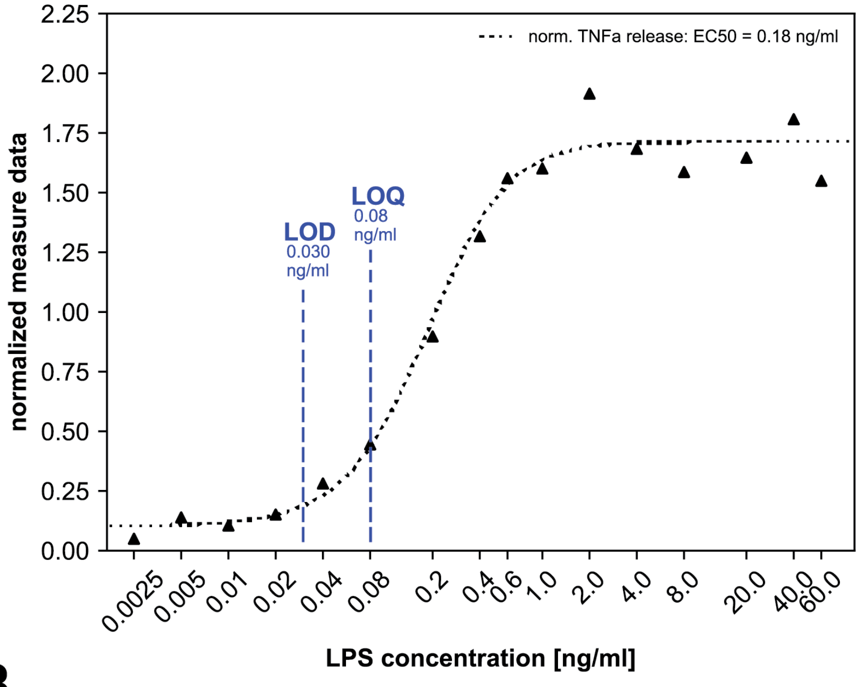


**B**

| feature | LOD Mean Activity | LOD (LPS) [ng/ml] | LOQ (LPS) [ng/ml] |
|---|---|---|---|
| TNFa release [pg/ml] | 250.44 pg/ml | 0.03 | 0.08 |

***Figure S5:*** **Mimicking endotoxin contamination on SiO2-OH ENM with LPS.** To mimic endotoxin contamination on ENM, different doses of LPS were added to the PBMCs and activated with a fixed amount (100 µg/ml) of $SiO_2$-OH (25 nm) NP that was shown to not induce general cytotoxicity in PBMCs. TNFα release was analyzed from obtained HTRF data. Limit of Detection (LOD) and Limit of Quantification (LOQ) were calculated by determining the mean activity of the blank (no LPS, no NP), with LOD_activity=mean+3std and LOQ_activity=mean+10std [1]. LPS concentrations of LOD and LOQ were calculated at the corresponding activity levels through the curve fit parameters of the four parametric logistic regression.
